# Energy predicts global mountain endemic plant richness better than environmental heterogeneity

**DOI:** 10.1101/2025.03.20.644469

**Authors:** Yuanyuan Gu, Jiaxin Ji, Zhimin Li, Zihan Jiang, Wenguang Sun

**Author notes:** **Corresponding author –**.

## Abstract

The drivers of biodiversity in mountain ecosystems have long been a central focus in ecologists. Increasing evidence suggests that energy is a key determinant of mountain species diversity; however, whether this pattern holds universally across different mountain ecosystems remains unclear, especially as there may be differences between different plant taxa. To address this knowledge gap, we selected mountain endemic plant genera from global biodiversity hotspots to explore the main drivers of diversity of different taxa in the mountains. Our results indicate that energy is the key driver of endemic plant richness in mountain regions worldwide, particularly for endemic tree and shrub taxa, while endemic herb richness is shaped by both energy and environmental heterogeneity. Regional studies have shown that energy availability drives total endemic plant groups in 70% of mountain regions. Specifically, energy is the dominant driver for 86% of endemic tree groups and 67% of endemic shrub groups, whereas endemic herb groups are the least influenced by energy, with only 50% of mountain regions showing energy as the primary driver. Our findings indicate that energy availability is the predominant factor shaping the diversity of endemic plant groups in mountain ecosystems worldwide. Therefore, mountain ecological conservation efforts should focus extensively on energy input aspects.

**Highlights:**

- Climate energy is the main driver of the richness of montane endemic plant taxa, especially for tree and shrub, whereas herb richness is determined by a combination of energy and environmental heterogeneity.
- Environmental heterogeneity predominantly drives endemic taxa richness in the Cape of Good Hope, whereas both climatic energy and environmental heterogeneity jointly influence endemic taxa richness in the Andes and Japan. In all other mountain ranges, climatic energy is the primary determinant.
- Endemic tree taxa have the highest number of mountains dominated by climatic energy, followed by endemic shrub taxa, and endemic herb taxa are more affected by environmental heterogeneity.

## 1. Introduction

Globally, mountains are recognized as biodiversity hotspots(Testolin et al., 2021). which not only support rich species and endemic biotic communities but also play an important role in maintaining ecological balance(Rahbek et al., 2019). Mountain ecosystems provide diverse habitats for species as well as mountain microenvironments, and are the most biodiverse terrestrial units on Earth(Zu and Wang, 2022). Energy has been increasingly recognized as a key driver of mountain species diversity(Francis and Currie, 2003; Bouchard et al., 2024), however, whether it universally dominates across different mountain ecosystems remains uncertain, especially as it may vary among plant taxa.

The collision of tectonic plates millions of years ago led to the uplift of the Earth’s surface. During this uplift, as the terrain became more complex and ecological conditions more diverse, many species underwent significant differentiation in their adaptation to new environments (Noori et al., 2024). In particular, mountain uplift facilitated the emergence of small-scale species divisions and drove extensive speciation (Ding et al., 2020). The terrain features resulting from surface uplift, such as valleys, cliffs, and river valleys, not only increased environmental heterogeneity but also created new ecological niches (Liu et al., 2019). Due to the complexity of microclimates and topographic barriers in mountainous regions, gene flow between different populations is restricted, intensifying population isolation (Myster, 2021). Isolated populations then evolve into new species as they adapt to different environmental conditions. In high mountain terrains, the windward and leeward slopes exhibit distinct ecological characteristics. The windward slopes, receiving abundant precipitation and stable sunlight, provide high energy inputs, creating optimal growth conditions for organisms, which results in higher species richness in these areas (Campbell et al., 2010; Lima et al., 2012; Cao et al., 2024). In contrast, the leeward slopes are relatively dry, with more harsh conditions for water and light, and this ecological pressure likely drives species to undergo adaptive evolution, resulting in the formation of endemic species with unique adaptations (Mulch, 2016; Spicer et al., 2020). Moreover, the leeward slopes may exert selective pressures on certain species, promoting their adaptive evolution under specialized environmental conditions. By comparing the ecological differences between windward and leeward slopes, we can gain deeper insights into the profound impacts of environmental change on species distribution and the evolution of diversity.

Endemic taxa are ideal subjects for studying these hypotheses because their evolutionary histories are confined to specific geographic regions and are typically unaffected by the spread of exotic species or gene flow. As a result, the evolutionary and ecological characteristics of these taxes can more purely reflect the influence of local environmental factors on species adaptation and diversity. In studying these taxa, we can more accurately observe the effects of environmental and topographic factors on species evolutionary processes, avoiding the confounding effects of external gene flow on species genetic background and ecological adaptation (Kruckeberg and Rabinowitz, 1985). These taxes are influenced by both energy input, which affects ecosystem productivity, and environmental heterogeneity, which drives habitat diversity. Energy input impacts plant growth and reproductive potential, thereby affecting species distribution and adaptive traits (Hall et al., 1992), while environmental heterogeneity promotes evolutionary adaptations by providing diverse habitats and selective pressures for plants (Anderson et al., 2014). However, the response mechanisms of endemic plant taxa are not uniform; instead, they vary significantly depending on the biological characteristics, ecological niche requirements, and evolutionary histories of the taxa (Mayor et al., 2017; Feldman et al., 2024). Therefore, understanding the response mechanisms of these taxes with-in specific montane ecosystems not only aids in elucidating the roles of energy and environmental factors in species distribution but also provides crucial insights into the dynamic changes of ecosystems. Based on the differences between the energy and environmental heterogeneity hypotheses outlined above, we propose the following hypotheses: (i) Mountains primarily influenced by energy factors include the Rocky Mountains and surrounding ranges in North America, the Appalachian Mountains, the European Alps and surrounding ranges, other mountains in Africa, the western highlands of Asia, the Himalayas-Tibetan Plateau-Hengduan Mountains, and Japan. (ii) Mountains primarily influenced by topographic factors include the Cape of Good Hope region in South Africa. (iii) Mountains most likely influenced by both energy and topographic factors include the Andes in South America.

This study aims to investigate the differential effects of energy input and environmental heterogeneity on various taxa, with a particular focus on the distribution and adaptive mechanisms of endemic plant genera in montane ecosystems worldwide. Through this research, we seek to elucidate the role of these key factors in shaping the diversity of endemic plants, thereby addressing existing knowledge gaps in the field. The core of the study lies in advancing our understanding of the mechanisms underlying montane biodiversity formation, thus providing theoretical support for the conservation and management of global montane ecosystems. Additionally, the findings will offer scientific insights for predicting biodiversity dynamics in the context of future environmental changes.

## 2. Materials and methods

### 2.1. Species data

We extracted scientific names of endemic mountain plants from published articles and then verified them using Plants of the World Online (https://powo.science.kew.org/), Global Biodiversity Information Facility (GBIF, https://ww-w.gbif.org/), and The World Flora Online (https://www.worldfloraonline.org/). We defined endemic genera as those where the ratio of endemic species to total species (endemic species/total species) is ≥0.75 (Dagnino et al., 2020), resulting in 722 endemic genera. We obtained species distribution data for these endemic genera from the Global Biodiversity Information Facility (GBIF, https://www.gbif.org/) and used the R package ‘*Coordinate Cleaner*’ to clean the data. Records lacking coordinate information were excluded, and abnormal coordinates, zero-value coordinates, duplicate latitude-longitude pairs, and other invalid geographical records were identified and removed to ensure data accuracy and reliability (Zizka et al., 2019). A total of 202,201 occurrence records were retained for analysis.

We classified endemic genera into tree, shrub, and herb based on their taxonomy in Plants of the World Online (https://powo.science.kew.org/), Global Biodiversity Information Facility (GBIF, https://www.gbif.org/), The World Flora Online (https://www.worldfloraon-line.org/), and China Plant Database (https://www.iplant.cn/) or their descriptions in plant specimens. The species distribution data were then mapped onto a 0.25° × 0.25° grid system (as shown in Figure 2), with grid cells covering less than 50% of the total area excluded. In calculating species richness, the number of endemic species within each grid was used as the species richness metric for that grid.

### 2.2. Parameter selection and calculation

We selected eight parameters to describe energy and topography from different perspectives (Table 1). Four energy parameters include: Mean annual temperature (MAT), Mean annual precipitation (MAP), Actual evapotranspiration (AET), and Mean annual solar radiation (MASR) (O’Brien, 1998; Clarke and Gaston, 2006; Yang et al., 2016; Coelho et al., 2023). MAT reflects the thermal input of the climate, directly influencing the growing season and the intensity of biological activity in plants, making it a fundamental parameter for measuring energy input. MAP, together with temperature, determines the availability of water, which is crucial for plant growth and ecosystem productivity, thus indirectly affecting the availability of energy. AET represents the exchange between water and energy; higher AET typically indicates efficient use of both water and energy, promoting ecosystem productivity. MASR is the primary driver of photosynthesis in plants and a key variable in ecosystem energy flow. Four topographic parameters include: Mean elevation (ME), Terrain undulation (TU), Standard deviation of elevation (STDE), and the ratio of surface area (TRSA). The overall elevation of mountain regions directly affects climate conditions; thus, ME captures this influence.

**Table 1.** Parameter selection.

| Argument | Unit | Abbreviations | References |
| --- | --- | --- | --- |
| Mean annual temperature | °C | MAT | (Fang et al., 2004) |
| Mean annual precipitation | mm | MAP | - |
| Actual evapotranspiration | mm | AET | (Yang et al., 2016) |
| Mean annual solar radiation | j | MASR | - |
| Mean elevation | m | ME | (Yu et al., 2018) |
| Terrain undulation | m | TU | - |
| Standard deviation of elevation | m | STDE | - |
| The ratio of surface area | - | TRSA | - |

Higher terrain undulation (TU) and larger standard deviation of elevation (STDE) provide more ecological niches and biodiversity habitats, while TRSA reflects the topographic roughness of a region (Fang et al., 2004; Yu et al., 2018). All climate data were sourced from WorldClim (https://www.worldclim.org/), covering the period 1970-2000 with a spatial resolution of 30 arc-seconds. Elevation data were downloaded from the GDEM V3 30 arc second Digital Elevation Model (https://www.gscloud.cn/). The global mountain polygon data were sourced from GMBA mountain inventory_V1.2 (Körner C, 2017). All variables were aggregated to a 0.25° × 0.25° grid cell. MAT and MAP were calculated for each grid using the ArcGIS 10.8 (Esri, 2020) Zonal Statistics tool. AET was calculated based on the formula using MAT and MAP (Yang et al., 2016), MASR was derived by calculating the monthly mean solar radiation using the ArcGIS 10.8 Zonal Statistics tool, followed by the calculation of the annual mean. The downloaded elevation data were mosaicked using the “Mosaic to New Raster” tool in ArcGIS 10.8, and the Zonal Statistics tool was then applied to calculate ME, TU, and STDE. TRSA was calculated using the DEM Surface Tool (Jenness, 2013) downloaded from https://www.jennessent.com/arcgis/surface_ar-ea.htm, followed by the use of the Zonal Statistics tool.

### 2.3. Statistical analysis

We first constructed univariate generalized additive models (GAMs), as GAMs can accommodate both linear and nonlinear relationships (Gomez-Rubio, 2018), making them well-suited for capturing the complex associations commonly observed in ecological studies. Based on the characteristics of our data, we selected either a Poisson or quasi-Poisson distribution to ensure model adaptability and optimal fit. By comparing model significance, AIC, or QAIC values, we identified the models with stronger explanatory power, ensuring the reliability and accuracy of our results (Burnham, 2002).

Prior to model selection, we first addressed multicollinearity among the variables. When a strong correlation was detected between two variables (Pearson’s |r| > 0.7), we prioritized removing the variable that showed a weaker independent relationship with species richness (i.e., the variable with a higher AIC or QAIC value). This approach effectively mitigates the impact of multicollinearity on model outcomes, thereby enhancing the model’s explanatory power and reliability.

Model selection was performed using the dredge function from the R package ‘*MuMIN’* (Barton, 2012). In the analysis, we explored various combinations of the selected variables fitted generalized additive models (GAMs) with Poisson or quasi-Poisson error distributions. The generated models were then evaluated and ranked based on the Akaike Information Criterion (AIC) or its corrected version, QAIC. To further assess the importance of each variable, we calculated its cumulative Akaike weight, *wi*, across all models, thereby determining its relative contribution (Letten et al., 2013).

## 3. Results

### 3.1. Overall richness pattern of endemic taxa

Our results show that endemic taxa are more diverse in the Andes of South America and the Cape of Good Hope region of South Africa compared to other mountain regions (Fig 1). This study examines the influence of energy factors and environmental heterogeneity on the diversity of endemic plant taxa across global mountain ecosystems. The findings indicate that energy factors, particularly climatic variables, play a dominant role in shaping the richness of total endemic plant taxa (Fig 2: All), with significant nonlinear relationships observed between these factors and species richness. For endemic tree and shrub taxa, key energy factors, including MAT, MAP, AET, and MASR—also exert a primary influence on richness (Fig 2: Tree and Shrub). In contrast, the richness of endemic herbaceous taxa is shaped by the combined effects of energy and topographic factors (Fig 2: Herb), highlighting the critical role of their interaction in determining herbaceous plant distribution patterns.

**Fig. 1.**
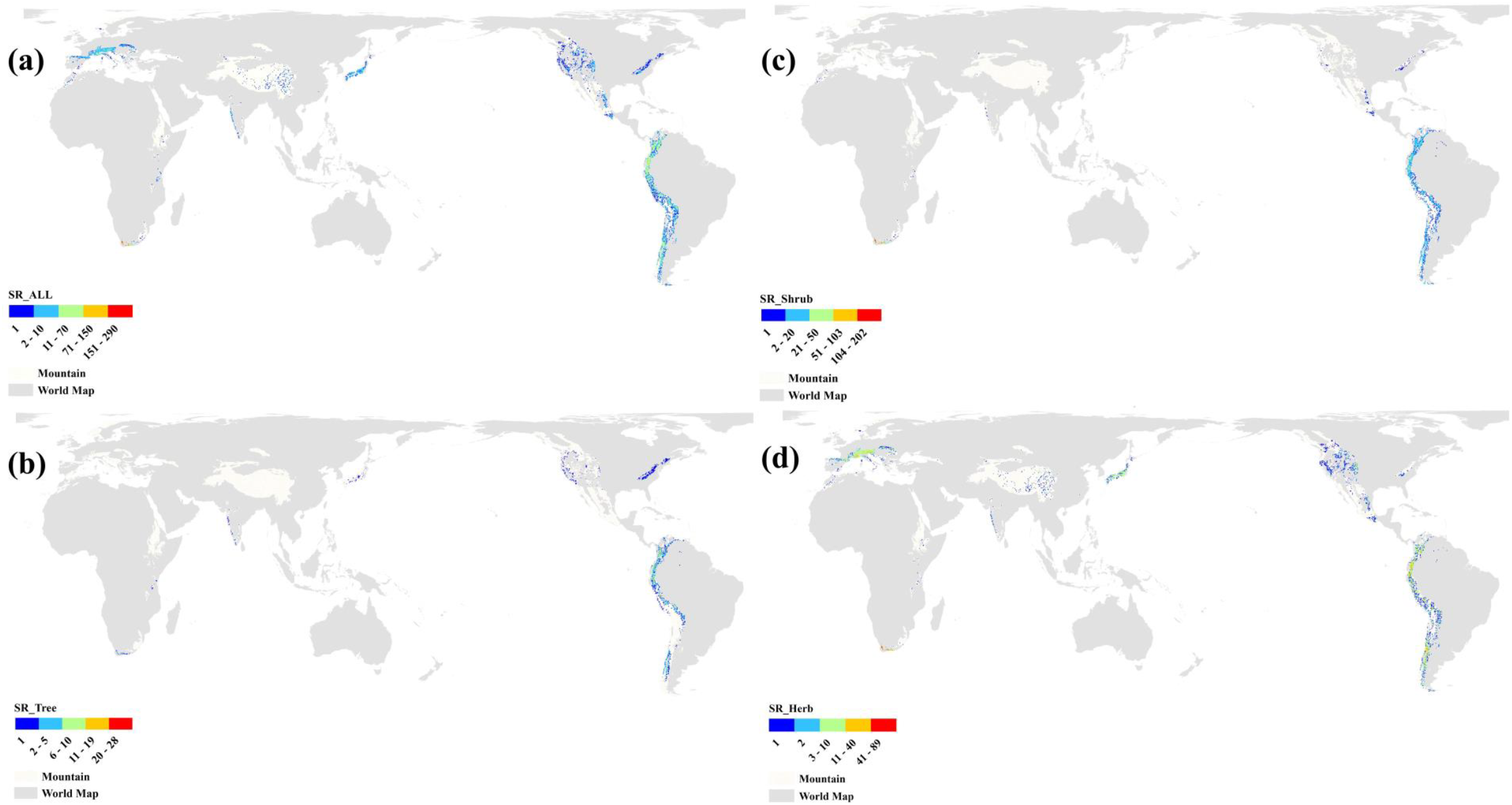
Diversity patterns of endemic taxa (a: all species; b: trees; c: shrubs; d: herbs) based on 0.25° × 0.25° grid cells.

**Fig. 2.**
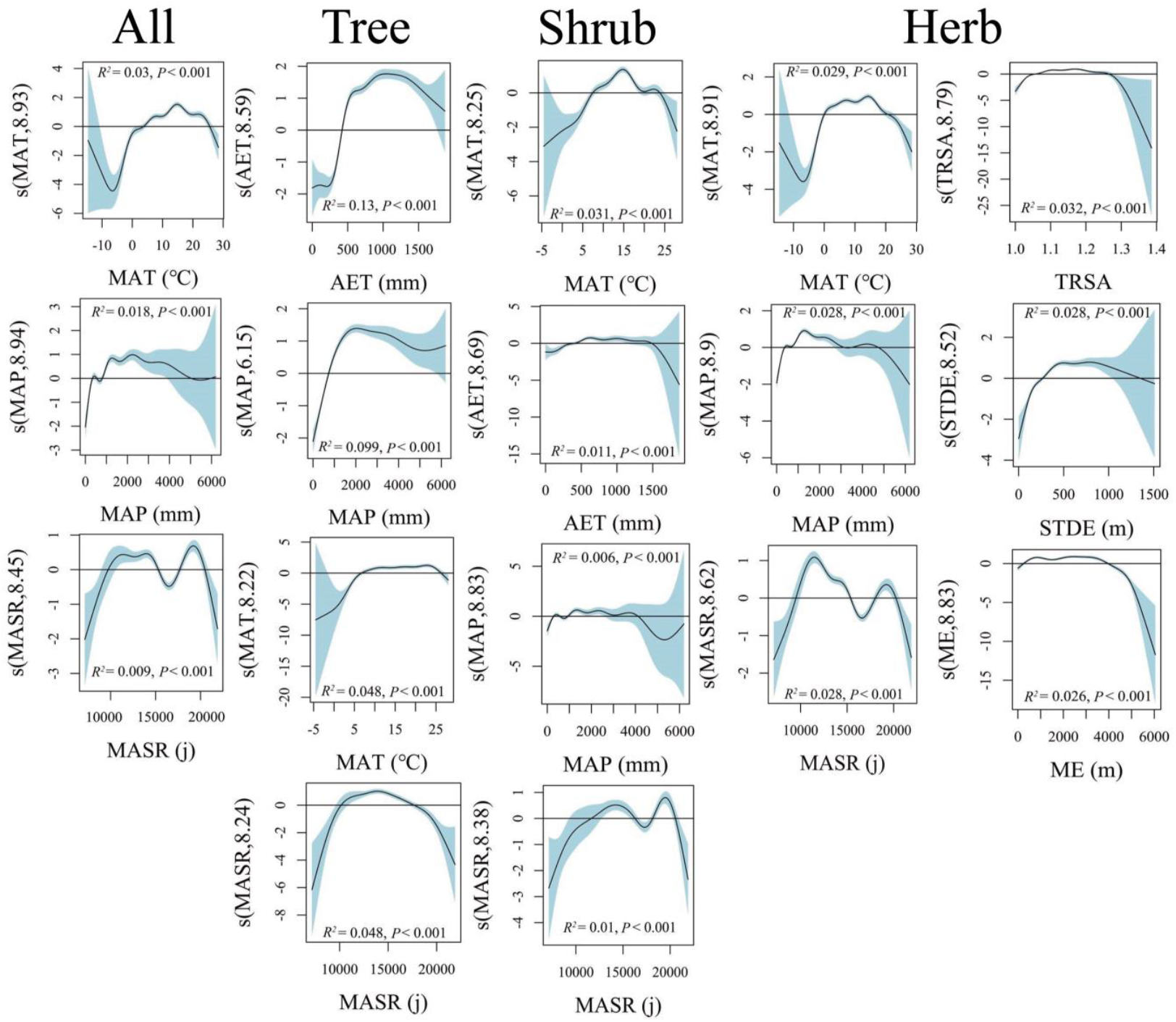
The variable factors that play a dominant role in the richness of total species, trees, shrubs, and herbs. Abbreviations are defined in Table 1.

Overall, energy input, particularly climatic energy, plays a pivotal role in shaping montane plant diversity. The diversity of different taxa is influenced by the interaction between energy and environmental heterogeneity. Future research should further investigate this complex relationship to enhance our understanding of its role in ecosystem dynamics.

### 3.2. Richness patterns of different endemic taxa in each mountain range

This study demonstrates that in most mountain regions, the richness of total endemic species is primarily driven by energy factors (Fig 3: All). In particular, climatic variables— MAT, MAP, AET and MASR—play a significant role in shaping species distribution and richness in the Rocky Mountains of North America, the Appalachian Mountains, the European Alps, and the Tibetan Plateau-Himalayas-Hengduan Mountains. These factors provide resources and habitats, thereby driving variations in species richness across these regions. However, in the Cape of Good Hope region of Africa, topographic factors—including TU, TRSA and ME—exert a dominant influence, significantly affecting the richness of endemic taxa. The complex topographic conditions and alpine climate create diverse habitats, promoting high levels of species differentiation and local adaptation, ultimately enhancing biodiversity. Additionally, we identified distinct patterns in the Andes of South America and Japan, where both energy and topographic factors jointly influence endemic species richness. In the Andes, species diversity is shaped not only by climatic energy but also by the constraints imposed by topographic features, reflecting the interplay between terrain complexity and climate. This finding suggests that in certain regions, the combined effects of energy and topography may have profound implications for species distribution and biodiversity patterns, warranting further investigation into their interactions across spatial scales.

**Fig. 3.**
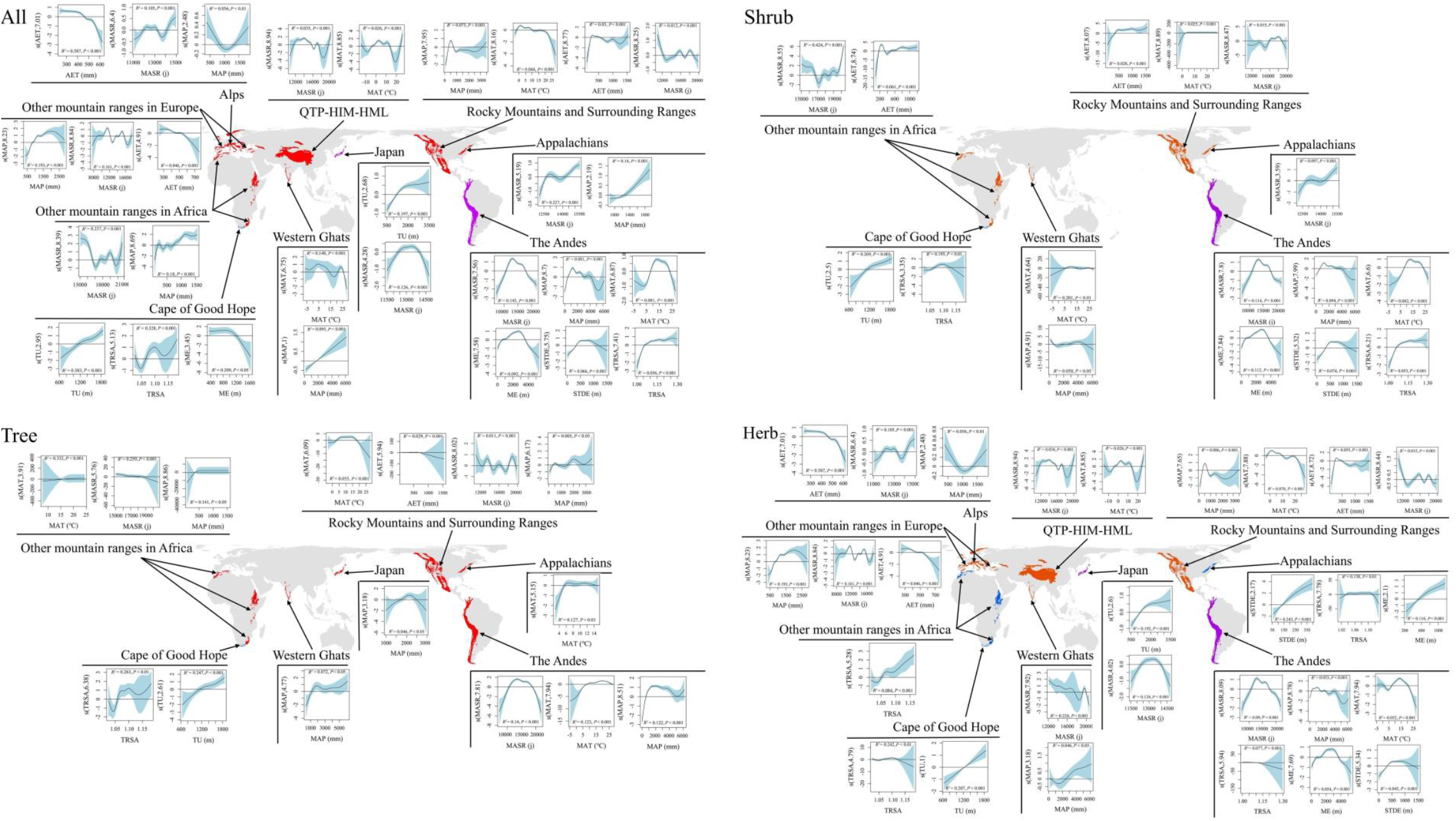
Factors that dominate different taxes in each mountain range. Red is energy dominance, blue is topography dominance, and purple is codominance. Abbreviations are defined in Table 1.

From the perspective of endemic tree taxa (Fig 3: Tree), topographic factors remain the dominant drivers in the Cape of Good Hope region of Africa, particularly in regions with complex alpine terrain and extreme climatic conditions, where these features significantly constrain the expansion of tree taxa. However, in other mountain regions, energy factors— particularly climatic energy—play a primary role, with energy input directly influencing the species richness of endemic trees in these areas.

For the richness of endemic shrub taxa (Fig 3: Shrub), topographic factors remain the primary drivers in the Cape of Good Hope region of Africa, where the region’s complex terrain constrains the expansion of endemic shrubs and promotes population isolation. In the Andes of South America, both energy and topographic factors jointly shape shrub diversity, indicating a strong interplay between climatic and geological influences. In most other mountain regions, however, energy factors serve as the dominant forces driving variations in species diversity.

For the richness of endemic herb taxa (Fig 3: Herb), topographic factors play a prominent role in the Cape of Good Hope region of Africa, as well as in mountain ranges outside this region and in the Appalachian Mountains of North America. The complex terrain and localized climatic conditions in these areas create unique habitats, leading to high species richness of endemic herbaceous plants. However, in most other mountain regions, energy factors remain the dominant drivers, particularly in the Alps and the Rocky Mountains of North America. Notably, in the Andes of South America and Japan, the richness of endemic herbaceous taxa is shaped by the combined influence of both energy and topographic factors, highlighting the critical role of their interaction in determining herbaceous diversity in these specific regions.

In summary, the influence of energy and topographic factors on the diversity of endemic plant taxa varies significantly across mountain regions. In most mountain ranges, energy factors primarily drive species richness, whereas in certain regions with extreme climates and complex terrain, topographic factors exert a more pronounced influence. Additionally, in some areas, the combined effects of energy and topography jointly shape the diversity patterns of endemic plant taxa, highlighting the intricate interplay between these factors in montane ecosystems.

## 4. Discussion

### 4.1. Preface review and summary of research results

For a long time, ecologists and biogeographers have argued that environmental heterogeneity (such as climate extremes, topographic barriers, etc.) is the primary driver of high biodiversity in montane ecosystems (Dornelas et al., 2009; Rahbek et al., 2019). This view is based on the idea that species in mountain ecosystems are often subject to complex and fluctuating environmental pressures, which in turn promote species differentiation and adaptive evolution (Stein et al., 2014). However, recent studies have increasingly suggested that, in addition to environmental heterogeneity, energy input—particularly climate energy—may also play a critical role in the formation of mountain biodiversity (Tito et al., 2020; Wang et al., 2022). Adequate climate energy can enhance ecosystem productivity, thereby providing more abundant resources and habitats for species, which in turn fosters greater species diversity (Evans et al., 2005).

This study analyzes 722 endemic plant genera across global mountain ecosystems, revealing a significant relationship between energy, particularly climate energy—and mountain biodiversity. The results indicate that climate energy input plays a more prominent role in shaping species diversity in most mountain ranges, with its influence being more pronounced in certain regions compared to traditional environmental heterogeneity factors. This finding challenges the traditionally dominant view that environmental heterogeneity is the primary driver of biodiversity, offering a new perspective on understanding the diversity of montane ecosystems.

### 4.2. Comparison and explanation of results with existing theories

The results of this study are consistent with several regional studies, particularly those conducted in the Tibetan Plateau-Himalayas -Hengduan Mountains region. Existing research has indicated that the monsoon, as a key climatic factor, significantly influences species differentiation and diversity in this area (Ding et al., 2020). This study highlights the role of climatic factors, particularly climate energy and the monsoon, in shaping species distribution and diversity within montane ecosystems, further validating the central role of climate conditions in biodiversity formation. Our findings also reveal a significant impact of climate energy—especially the monsoon—on species diversity across several mountain regions, supporting the notion that climatic factors, particularly energy input, are crucial drivers of montane plant diversity. Specifically, climatic factors such as solar radiation and precipitation directly influence plant growth, distribution, and community structure (Bouchard et al., 2024). Sufficient energy promotes plant reproduction and growth (Wright, 1993; Surangi et al., 2007), thereby affecting community diversity and species composition. Therefore, energy input is a critical determinant of plant community structure and ecological processes.

### 4.3. Differences in plant groups’ perception of energy requirements and environmental inhibitions

Different plant taxa have varying demands for photosynthesis, temperature, and moisture, which results in heterogeneous effects of energy on them(Francis and Currie, 2003). The response of different plant taxa to energy and environmental heterogeneity also varies, with the relative importance of these factors differing across taxa (Aparicio et al., 2008; Maes et al., 2020). For instance, herbaceous plants are generally more sensitive to environmental changes, and topographic barriers more readily lead to population isolation, thereby driving speciation(Morente-López et al., 2018; Li et al., 2022). In contrast, woody plants are relatively more resilient to environmental pressures and are more influenced by energy factors, such as light and climate energy, making their distribution and diversity more dependent on energy availability (Chelli-Chaabouni, 2014).

### 4.4. The complexity of environmental heterogeneity

Although the relationship between energy and species diversity is significant in most cases, this study does not entirely exclude the importance of environmental heterogeneity. In certain mountain regions, due to unique environmental conditions such as alpine climates and soil types, environmental heterogeneity may still lead to the isolation and differentiation of distinct biotic communities. Therefore, environmental factors continue to play a crucial role in specific areas, particularly in regions with extreme topography and climate, where the complexity and diversity of the environment have profound effects on species distribution and diversity (Stein et al., 2014; Deák et al., 2021; Barczyk et al., 2023).

### 4.5. Interaction of energy and environmental heterogeneity

While energy is a key factor in shaping biodiversity, environmental heterogeneity still plays a significant role under specific conditions, such as complex topography and extreme climates (Dufour et al., 2006). In montane ecosystems, energy and environmental heterogeneity do not act in isolation but are intertwined, jointly influencing species distribution and diversity (Antonelli and Sanmartín, 2011; Trew and Maclean, 2021). Energy impacts ecosystem productivity, providing the resources necessary for species, while environmental heterogeneity further promotes species differentiation and evolution by creating diverse habitats and selective pressures (Hughes and Eastwood, 2006; Bouchenak-Khelladi et al., 2015). Therefore, in mountain ecosystems, the interaction between energy and environmental heterogeneity must be considered comprehensively to fully understand the mechanisms underlying biodiversity formation.

The effects of energy and environmental heterogeneity are not the result of a single factor, but rather the interaction between the two across different temporal and spatial scales. For instance, under certain seasons or specific climatic conditions, adequate energy may be the dominant factor determining species distribution (Tonkin et al., 2017; Arenas-Navarro et al., 2019), as it directly influences plant growth and reproduction. However, under other conditions, such as extreme climatic events or in areas with complex topography, environmental heterogeneity may become the key factor limiting species distribution and diversity (Deák et al., 2021). Therefore, the interaction between energy and environmental heterogeneity needs to be analyzed comprehensively across various temporal and spatial scales to more accurately reveal their joint impact on ecosystem structure and species diversity.

### 4.6. Limitations and future directions of the study

This study primarily relies on global mountain data, which, while revealing the overall influence of energy and environmental factors on plant diversity, does not fully account for the subtle differences in local ecosystems. This limitation may have led to the oversight of certain local environmental variables, such as soil properties and wind direction, which could play an important role in species distribution and diversity in specific regions (Moeslund et al., 2013; Delgado-Baquerizo et al., 2020; Dai et al., 2023). Future research could delve deeper into the responses of different plant taxa to energy and environmental heterogeneity factors using higher-precision local data, particularly in the context of climate change, to better understand how these factors influence montane plant diversity and ecological processes. In future studies, we will further clarify the relative importance of energy and environmental heterogeneity in shaping global montane plant diversity by comparing plant diversity data across different mountain ranges and climatic zones. By refining the ecological characteristics of different regions, we can more clearly reveal how energy and environmental factors interact across various temporal and spatial scales, affecting species distribution and community structure.

## 5. Conclusion

This study primarily relies on global mountain data, which, while revealing the overall influence of energy and environmental factors on plant diversity, does not fully account for the subtle differences in local ecosystems. This limitation may have led to the oversight of certain local environmental variables, such as soil properties and wind direction, which could play an important role in species distribution and diversity in specific regions (Moeslund et al., 2013; Delgado-Baquerizo et al., 2020; Dai et al., 2023). Future research could delve deeper into the responses of different plant taxa to energy and environmental heterogeneity factors using higher-precision local data, particularly in the context of climate change, to better understand how these factors influence montane plant diversity and ecological processes. In future studies, we will further clarify the relative importance of energy and environmental heterogeneity in shaping global montane plant diversity by comparing plant diversity data across different mountain ranges and climatic zones. By refining the ecological characteristics of different regions, we can more clearly reveal how energy and environmental factors interact across various temporal and spatial scales, affecting species distribution and community structure.

## Declaration of competing interest

The authors declare that they have no known competing financial interests or personal relationships that could have appeared to influence the work reported in this paper.

## Data statement

When an article is accepted, we will upload the data to Science Data Bank.

## Notes

### Competing Interest Statement

The authors have declared no competing interest.

